# The vagus nerve promotes memory via septo-hippocampal acetylcholine: Implications for obesity-induced cognitive dysfunction

**DOI:** 10.1101/2025.06.11.659206

**Authors:** Logan Tierno Lauer, Léa Décarie-Spain, Anna M.R. Hayes, Andrea N. Suarez, Alexander Bashaw, Molly E. Klug, Alicia E. Kao, Robert Cheng, Jessica J. Rea, Keshav S. Subramanian, Anna Nourbash, Kristen N. Donohue, Lindsey A. Schier, Kevin Myers, Scott E. Kanoski

## Abstract

The vagus nerve relays critical metabolic information between the gastrointestinal (GI) tract and the brain. Recent findings highlight a role for vagus nerve-mediated gut-brain signaling in regulating higher-order cognitive processes, although the underlying mechanisms remain poorly understood. Here we demonstrate that nutrient consumption promotes hippocampal-dependent memory function via vagus nerve-mediated acetylcholine (ACh) release in the dorsal hippocampus (HPCd) from medial septum (MS) neurons. In vivo analyses reveal that HPCd ACh release is engaged during nutrient consumption, and that this response is abolished in animals that received MS cholinergic neuron ablation, subdiaphragmatic vagotomy (SDV), or early-life Western Diet (WD) maintenance. MS cholinergic neuron ablation, SDV, and early-life WD also impaired memory for meal location, suggesting that this gut-brain pathway functions to encode memories for eating events. Collectively, results identify a neurobiological mechanism whereby nutrient consumption promotes memory function, and reveal that disruption of this vagal-brain signaling system mediates WD-associated memory impairments.

## Introduction

Communication between the gastrointestinal (GI) tract and the brain is critical in maintaining food intake control and overall energy regulation ^1–4^. Emerging work also identifies a role for the gut-brain axis in higher-order cognitive functions ^5,6^. A key conduit of communication between the gut and the brain is the vagus nerve, whose afferent/sensory neurons (vagal afferent neurons, VAN) transmit a wide array of metabolic information from the GI tract to the central nervous system ^7^. Recent evidence indicates that VAN influences on brain processes extend beyond traditional brainstem targets to include transsynaptic communication to higher-order structures involved in cognition, including the hippocampus (HPC)^8–10^, a critical mediator of episodic and spatial navigation-based memory ^11–13^.

The medial septum (MS) is an anatomical relay of neural communication between the caudal brainstem, where GI VAN synapse, and the dorsal subregion of the hippocampus (HPCd)^10^. The MS contains a population of acetylcholine (ACh)-producing neurons, and septo-hippocampal cholinergic signaling promotes hippocampal plasticity and memory encoding ^14–16^. However, the upstream mechanisms that regulate septo-hippocampal memory systems are poorly understood. Here we posit that this memory-promoting circuitry is guided by visceral sensory pathways, including GI meal-associated signals transmitted to the brain via the VAN.

Consumption of a high-fat and sugar-enriched “Western diet” (WD) impairs HPC-dependent memory and associated neural processes, particularly when consumed during early life periods of development ^17–20^. Recent findings reveal that early life WD exposure compromises HPC function through dysregulated HPCd cholinergic signaling ^21^. In parallel, WD consumption and obesity are linked with blunted VAN signaling, as animals exposed to WD demonstrate reduced brainstem responsiveness to satiation signals that are transmitted through VAN ^22–26^. Furthermore, vagus nerve ablation through sub-diaphragmic vagotomy (SDV) or through sensory-specific cholecystokinin-conjugated saporin-mediated VAN ablation disrupts HPC-dependent memory function in rats, coupled with reductions in HPC markers of synaptic plasticity and neurogenesis (brain-derived neurotrophic factor and doublecortin, respectively) ^10^. Consistent with these findings, either gastric distension or intestinal nutrient infusion robustly enhances cerebral blood flow to the HPCd ^27,28^. Collectively, these findings highlight a putative functional connection between GI VAN and septo-hippocampal ACh signaling, and raise the possibility that obesity-associated cognitive deficits may stem, in part, from a breakdown in this gut-VAN-HPCd connection.

In the present study we elucidate how GI-originating VAN signaling modulates HPC function. Results identify a role for nutritive signals in driving HPC-dependent memory. Eating behavior dynamically increases HPC ACh release, and these levels remain elevated after a meal. These effects are eliminated by either VAN ablation or MS ACh neuron deletion, and are also negatively impacted by an early-life junk food diet. These data collectively define novel mechanisms revealing how and where gut-vagus-HPC communication promotes memory, and provide mechanistic insights into how interoceptive dysfunction may contribute to cognitive decline.

## Results

### The gut peptide CCK promotes hippocampal activity via septal cholinergic neurons

Cholecystokinin (CCK) is an intestinally-derived hormone that promotes satiation and meal termination in part through paracrine action on the vagus nerve in response to the presence of nutrients in the small intestines ^29^. The medial septum (MS) is an anatomical relay connecting the caudal brainstem (where gut-derived VAN converge) to the HPC ^10^, and thus the MS may communicate gut vagal signaling to the HPC. To explore this hypothesis, rats were injected with either a 192IgG saporin immunotoxin in the MS (to ablate MS cholinergic neurons) or a control saporin (Fig. 1A). Following surgical recovery, the animals received peripheral administration (intraperitoneal; IP) of either vehicle (saline) or CCK (3𝜇g/kg; a dose that requires the vagus nerve for food intake reduction^1^), and immunofluorescence analyses were conducted to quantify c-Fos protein expression in the dentate gyrus (DG) and CA3 subregions of the HPCd. Results revealed that, relative to vehicle treatment, IP CCK elevated c-Fos expression in the DG granular layer and in the dorsal CA3 (CA3d) pyramidal layer in animals who received MS control saporin treatment. However, MS 192IgG saporin injections eliminated IP CCK-induced HPCd c-Fos induction, with expression comparable to vehicle treatments (Figure 1a-c, Supp. Figure 1a-c). To further corroborate that MS ACh transmission is engaged in the HPCd by peripheral CCK, rats were injected with an pAAV.hSynap.iAChSnFR and an optic fiber cannula in the HPCd (DG) to detect fluctuations in ACh release *in vivo* in awake and freely behaving animals (Figure 1e) that received either 192IgG saporin or control saporin in the MS (Figure 1d). Post surgical recovery, fiber photometry was used to measure HPCd ACh release in response to IP CCK and vehicle administration following an overnight fast (Figure 1d-f). Results revealed a sustained (∼10-40 min after injections) elevation in HPCd ACh release after IP CCK administration relative to vehicle administration in animals with MS control saporin treatment; an effect that was eliminated by ablation of cholinergic neurons in the MS (Figure 1g-i). To examine whether this effect is specific to vagally-mediated peptide signals, we next evaluated HPCd ACh release in response to IP vehicle vs amylin, a pancreatic hormone with food intake-reducing effects that do not require the vagus nerve^30^ . In contrast to CCK, amylin did not elicit any responses in HPCd ACh signaling relative to vehicle treatments (Supp. Figure 1d). Together, these data reveal that CCK, a gut-derived peptide signals that act through the vagus nerve, promotes HPCd ACh activity via MS cholinergic neurons.

**Figure 1:**
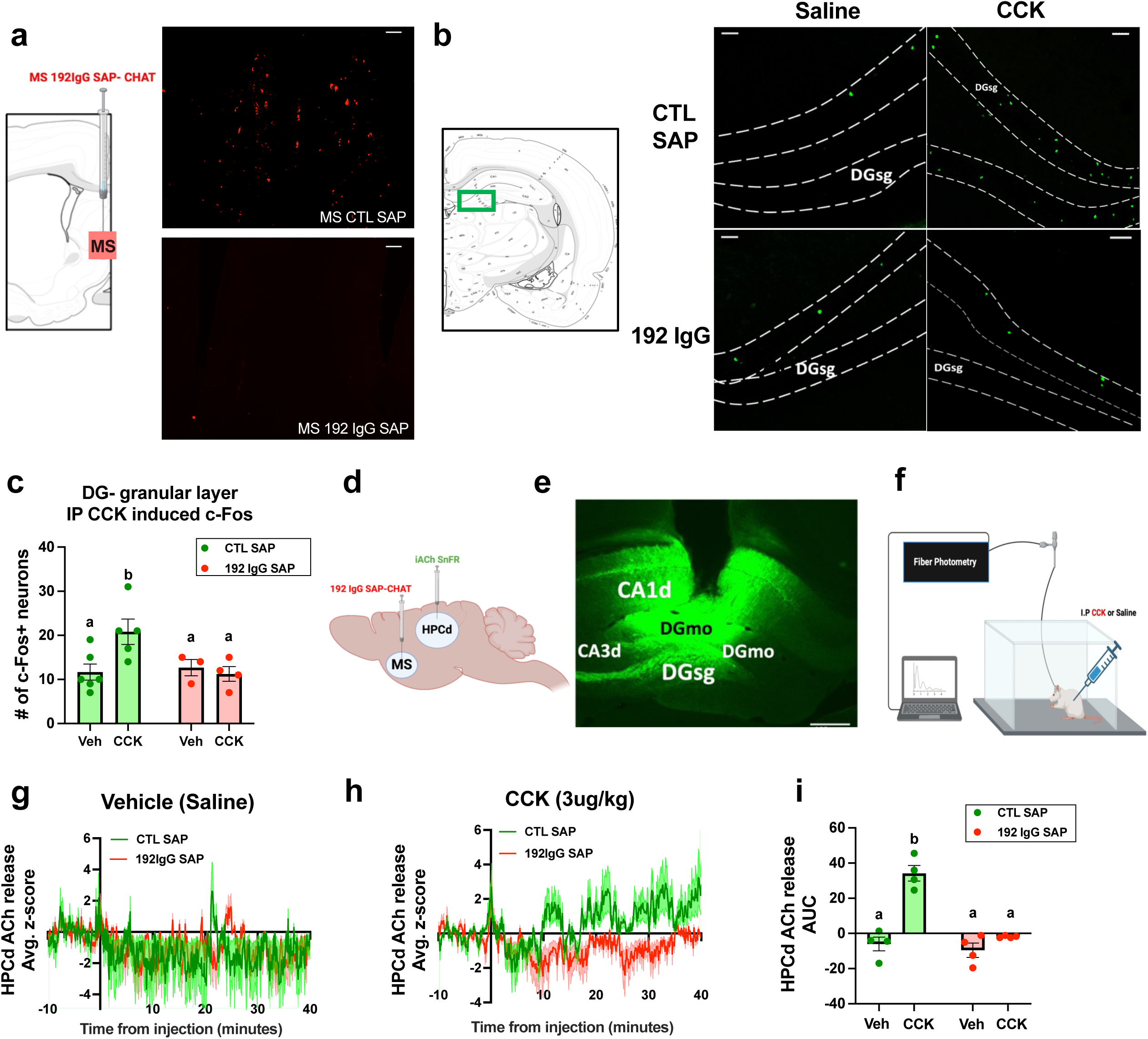
Peripheral CCK engages hippocampal activity via septal cholinergic neurons. **A.** Schematic and representative medial septum (MS) images of immunoreactive CHAT+ neurons in an animal with control saporin (top) and 192IgG saporin toxin (bottom). Scale bar = 30 microns. **B.** Schematic and representative images of dorsal dentate gyrus granular layer immunoreactive c-Fos+ neurons in control saporin and 192IgG saporin treated animals given IP CCK (3μg/kg) or vehicle (saline). Scale bar = 30 microns. **C.** Dorsal dentate gyrus granular layer c-Fos+ neurons quantification. (*n* = 6 Vehicle-CTL SAP; *n* = 4 CCK-CTL SAP; *n* = 3 Vehicle-192IgG sap; *n* = 4 CCK-192IgG SAP). Data were analyzed using two-way ANOVA with multiple comparisons. Treatment data with the same letter above are not significantly different. IP Treatment x saporin interaction *P =* 0.0066, CTL SAP Vehicle vs CCK \*\**P =* 0.0026; CCK treatment CTL SAP vs. 192IgG SAP \*\**P =* 0.0045. **D.** Schematic of surgical approach for 192IgG SAP in MS and iAChSnFR in HPCd. **E.** Representative HPCd image of GFP immunoreactive neurons containing iAChSnFR, scale bar = 100 microns. **F.** Schematic of fiber photometry recordings during IP CCK and vehicle treatments. **G.** Time-locked averaged HPCd ACh release to IP vehicle administration (*n =* 4 Vehicle-CTL SAP; *n* = 4 Vehicle-192IgG SAP). **H.** Time-locked averaged HPCd ACh release to IP CCK (3 μg/kg) administration (*n =* 4 CCK-CTL SAP; *n* = 4 CCK-192IgG SAP). **I.** HPCd ACh release AUC quantification in response to IP vehicle and CCK administration. Treatment data with the same letter above are not significantly different. Data were analyzed using two-way ANOVA with multiple comparisons; IP Treatment x saporin interaction \*\**P =* 0.0016; CTL SAP-Veh vs. CCK \*\*\**P =* 0.0006; CCK treatment CTL SAP vs. 192IgG SAP \*\*\**P =* 0.0008. Data are shown as mean ± SEM; \*\*\*\**P*<0.0001, \*\**P<*0.01.

### Meal consumption enhances hippocampus acetylcholine signaling

To evaluate HPCd ACh responses during physiological engagement of GI VAN, we recorded ACh release during meal consumption after an overnight fast. Timestamps were taken to analyze cholinergic activity during active eating periods (composed of eating bouts) vs. inter bout-periods (i.e., the periods between active eating bouts, during which animals are typically rearing or exploring) (Figure 2a). Results revealed that HPCd ACh release dynamically increased during active consumption of standard chow and subsequently decreases during inter bout-periods (representative trace from a single animal in Figure 2b; zoomed in trace shown in Suppl. Figure 2a-b; compiled group data in Figure 2c and 2d). Furthermore, HPCd ACh release was significantly elevated 5-min after voluntary meal termination (satiation state) compared to 5-min pre food consumption period (hunger state; Figure 2e-f). Interestingly, ΔHPCd ACh responses during eating bouts (but not during inter bout intervals) were significantly larger in magnitude during later consumption bouts (bouts occurring in the third tertile of a meal) in a meal compared to earlier ones (bouts occurring within the first tertile of a meal) (Supp. Figure 2c-d), suggesting that HPCd ACh release during eating tracks meal progression. Additionally, these food consumption-induced HPCd ACh responses were absent in animals that received 192IgG saporin in the MS relative to controls (Figure 2g-h), despite there being no differences in the amount of food consumed (Supp. Figure 2i). Meal period assessments in animals with control saporin reveal nonsignificant larger ACh responses during later consumptions bouts vs. earlier (Supp. Figure 2e-f). Collectively, these findings demonstrate that HPCd ACh release is highly responsive to naturalistic eating behavior, as demonstrated by its dynamic and sustained responses during active consumption periods and during satiated states. These results also further highlight the role of MS ACh neurons in HPCd responses to physiological engagement of the gastrointestinal tract.

**Figure 2:**
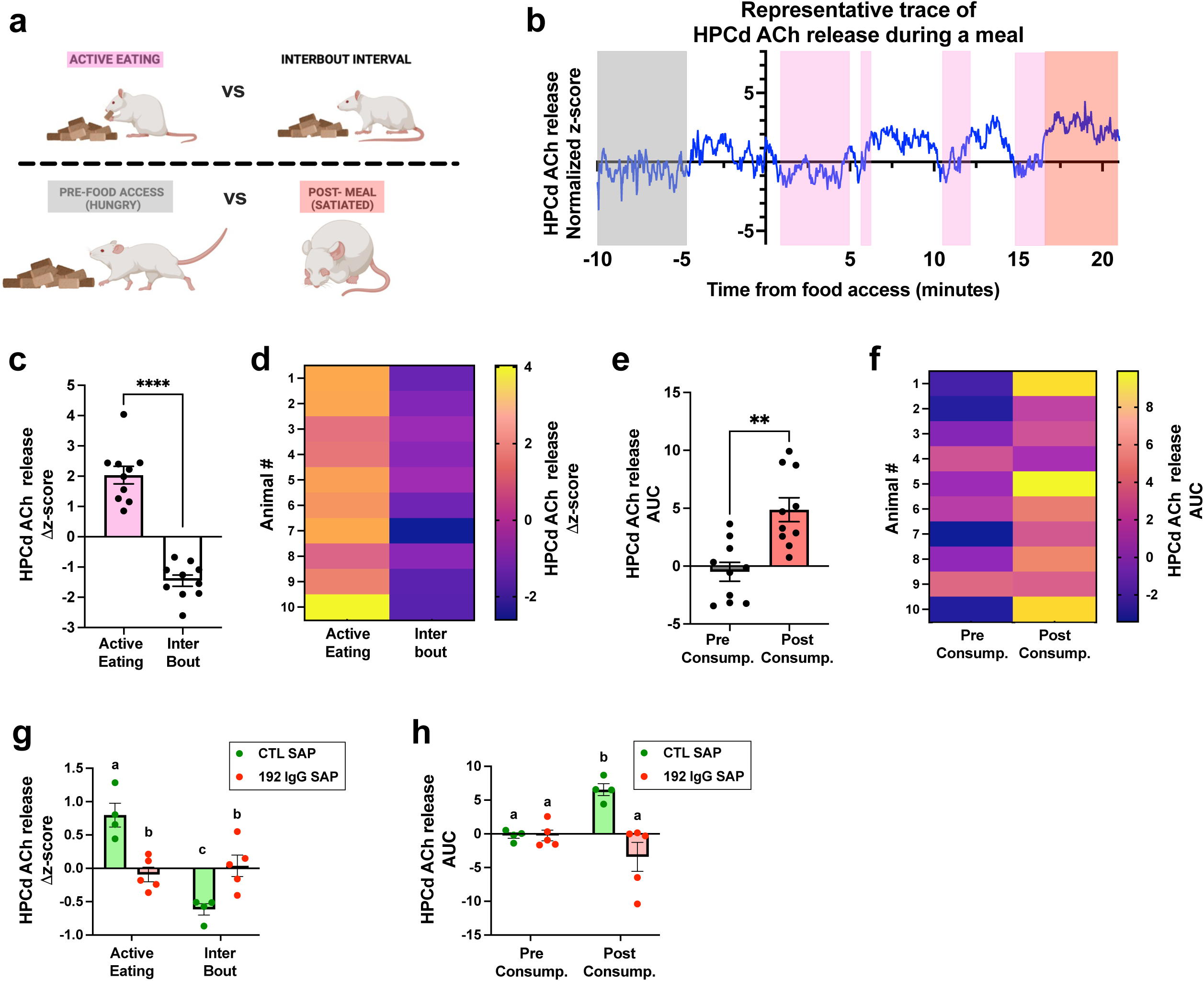
Meal consumption elevates septo-hippocampal cholinergic signaling. **A.** Schematic of events timestamped during fiber photometry recordings while animals consume a meal. **B.** Representative trace of HPCd ACh release during consumption of standard chow, time locked to moment of access to food. Pre-consumption period (fasted state) is marked in the gray box, active eating periods [pink boxes; quantified as the Δ in activity between start and end of eating bout], inter bout intervals [white boxes; quantified as the Δ in activity between the end of an eating bout and the beginning of the subsequent eating bout], and post-consumption period (satiated state) marked in the red box are represented. **C-F.** Fiber photometry recording of HPCd ACh release quantification during meal consumption (data analyzed using Student’s two-tailed paired *t*-test, *n* = 10 rats) with **C** change in HPCd ACh release during active eating periods [pink boxes] and inter bout intervals [white boxes], \*\*\*\**P* < 0.0001, **D** Heatmap representing each animal’s change in HPCd ACh release during active eating and inter bout periods. **E.** Area under curve (AUC) HPCd ACh release during 5-min pre-food consumption (fasted state; gray box) and 5-min post-food consumption (satiated state; red box), \*\**P* = 0.0054, and **F.** Each individual animal’s AUC values during the pre- and post-food consumption periods. **G-H.** Fiber photometry recordings of HPCd ACh release during food consumption in animals with either control saporin (CTL SAP; green), or 192IgG SAP (red) in the MS, all treatment data containing the same letter above are not significantly different; data were analyzed using two-way ANOVA with multiple comparisons, (*n*=4, CTL SAP; *n*=5, 192IgG SAP) with **G** depicting change in ACh release during active eating bouts and inter bout intervals for both experimental groups. Saporin treatment x eating period interaction \*\**P=*0.0027. CTL SAP, active eating vs. inter bout \*\*\**P*=0.0007; Active eating, CTL SAP vs. 192IgG SAP \*\**P*=0.0082; Inter bout, CTL SAP vs. 192IgG SAP \**P*=0.0325. **H.** HPCd ACh release AUC quantification during pre-consumption and post-consumption periods. Saporin treatment x energy state interaction *P=*0.053. CTL SAP, pre-consumption vs. post-consumption. \**P*=0.015; Post-consumption, CTL SAP vs. 192IgG SAP \*\**P*=0.0061. Data are shown as mean ± SEM; \*\*\*\**P*<0.0001, \*\**P<*0.01.

### Consumption-driven elevations in hippocampal acetylcholine responses require nutrients

HPCd ACh responses during meal consumption may be driven by the caloric consequences of the food, by specific macronutrients, by oral sensory stimuli (e.g., taste), and/or by the locomotive action of eating. To dissociate between these possibilities, we evaluated dynamic HPCd ACh responses during consumption of isolated macronutrients vs. no- and low-calorie sweetened solutions (Figure 3a). Results revealed that active consumption of an 11% sucrose solution induced the dynamic elevations in HPCd ACh release as previously observed with standard chow (Figure 3b). The sustained elevation in cholinergic tone after voluntary termination of the 11% sucrose solution relative to the pre-consumption hunger state was also observed (Figure 3b). Meal period analyses also revealed that ΔHPCd ACh release during later consumption bouts of 11% sucrose were significantly higher in magnitude than earlier ones, while there was no difference between the magnitude of ΔHPCd ACh release during early inter bout intervals vs. late ones (Supp. Figure 3a-b). Next, we assessed whether a calorically matched solution of 4.5% corn oil (pure lipid) would recapitulate these responses. Like carbohydrate consumption, lipid ingestion also elicited HPCd ACh responses during its active consumption periods, as well as a sustained elevation in ACh release after meal-termination (Figure 3c). ΔHPCd ACh responses were larger in magnitude during later consumptions bouts than earlier, but these analyses did not reach significance (Supp. Figure 3c-d). To interrogate whether these consumption-induced cholinergic responses were due to caloric content, we conducted the same experiment with 0.2% saccharin, which is an artificial sweetener that will engage sweet taste receptors but is devoid of calories. Results revealed no differences in ΔHPCd ACh release during active consumption of 0.2% saccharin vs. inter bout intervals, nor was there differential ACh release in the post-vs. pre-consumption period (Figure 3d). However, the animals consumed significantly less of the 0.2% saccharin solution compared to 11% sucrose or the 4.5% corn oil solutions (Figure 3f), and thus the different ACh response profiles may be based on volume of solution consumed and not caloric content. To control for ingestion volume while keeping caloric content very low, we repeated the experiment with a solution combining 0.2% saccharin with the addition of 2% sucrose, which resulted in consumption levels (volumes of solution) that were significantly higher than that of 11% sucrose and 4.5% corn oil (Figure 3f). Fiber photometry readings revealed no differences in ΔHPCd ACh release during active consumption of the 0.2% saccharin + 2% sucrose solution vs. inter bout intervals, and there was no difference in ACh tone between fasted and post-consumption states (Figure 3e). Further, there was no difference in the ΔHPCd ACh release during active consumption bouts, and inter bout intervals across meal period stages for either the 0.2% saccharin or the 0.2% saccharin + 2% sucrose solutions (Supp. Figure 3e-h). Together, these findings reveal that consumption-induced HPCd ACh engagement requires sufficient nutritive feedback.

**Figure 3:**
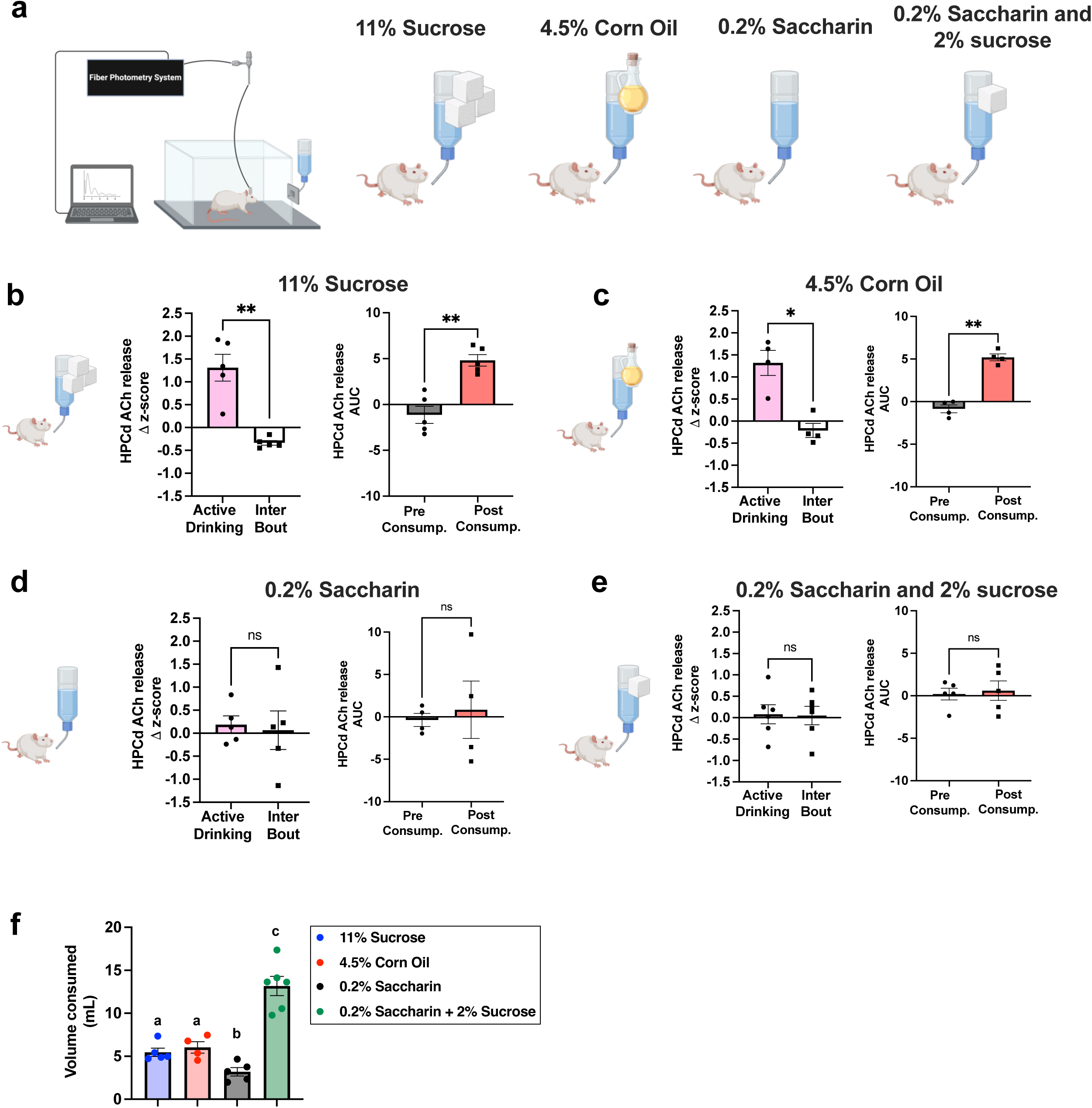
Meal-induced septo-hippocampal cholinergic signaling is driven by nutrients. **A.** Schematic of fiber photometry recordings of HPCd ACh release during consumption of select isolated macronutrients or artificial sweetener in solution. **B-E.** Fiber photometry recording of HPCd ACh release quantification during isolated macronutrient meal consumption (data analyzed using Student’s two-tailed paired *t*-test) with **B** reflecting the comparison between the change in HPCd ACh release during active eating periods vs. inter bout intervals during 11% sucrose consumption: *n*=5, \*\**P* =0.0032, as well as the AUC comparison between pre-consumption and post-consumption periods \*\**P*=0.0020. **C.** Change in HPCd ACh release during active eating periods vs. inter bout intervals during 4.5% corn oil consumption: *n*=4, \**P* =0.0406, as well as the AUC comparison between pre-consumption and post-consumption periods \*\**P*=0.0046. **D.** Change in HPCd ACh release during active eating periods vs. inter bout intervals during 0.2% saccharin consumption: *n*=5, as well as the AUC comparison between pre-consumption and post-consumption periods. **E.** Change in HPCd ACh release during active eating periods vs. inter bout intervals during 0.2% saccharin + 2% sucrose consumption: *n*=6, as well as the AUC comparison between pre-consumption and post-consumption periods. **F.** Amount of isolated macronutrient/sweetener solution that was consumed during fiber photometry recording sessions. All treatment data containing the same letter are not significantly different (11% sucrose vs. 0.2% saccharin \**P*=0.0223, 11% sucrose vs. 0.2% saccharin + 2% sucrose \*\*\**P*=0.0002, 4.5% corn oil vs. 0.2% saccharin \*\**P*=0.0096, 4.5% corn oil vs. 0.2% saccharin + 2% sucrose \*\**P*=0.0014, 0.2% saccharin vs. 0.2% saccharin + 2% sucrose \*\*\*\**P <* 0.0001. Data are shown as mean ± SEM; \*\*\*\**P*<0.0001, \*\**P<0.01*.

### Meal-induced elevations in hippocampal acetylcholine responses require the vagus nerve

Given that signals arising from gut macronutrient detection are transmitted via a variety of pathways ^31–33^ we next interrogated the role of the vagus nerve in meal-induced HPCd ACh responses in rats with either sham or sub-diaphragmatic vagotomy surgery (SDV) to disconnect GI vagus nerve signaling from the central nervous system. Elevations in HPCd ACh release during active consumption of chow, as well as the sustained elevation of cholinergic tone in the satiated vs fasted state were replicated in sham control rats. However, SDV abolished meal-induced ΔHPCd ACh release during active consumption, as well as the sustained elevation in cholinergic tone after voluntary meal termination without influencing the amount of food consumed (representative photometry traces in Figure 4a-b; group compiled data in Figure 4c-e). Meal period analyses of ΔHPCd ACh release in sham rats during active consumption revealed that cholinergic responses were higher in magnitude during later bouts compared to earlier ones only in sham controls, but not in SDV rats (Supp. Figure 4b-e). We next performed Western blot analyses in the HPC to quantify levels of proteins associated with ACh signaling, including choline acetyltransferase (ChAT), an enzyme that synthesizes ACh using choline and acetyl-CoA as precursors; vesicular acetylcholine transporter (VAChT), which carries ACh in vesicles for subsequent release upon synapse events, and acetylcholinesterase (AChE), an enzyme that degrades ACh after its release at the synaptic cleft. Results revealed that the SDV surgery significantly reduced VAChT levels in the HPCd compared to sham surgery (Figure 4g), with no group differences in ChAT or AChE expression (Figure 4f and h). These results suggest gut vagus nerve ablation induces a reduction in ACh vesicular transport in the HPCd, without eliciting any compensatory changes in the expression of other proteins involved in ACh neurotransmission. Overall, these data indicate that the vagus nerve is required for dynamic HPCd ACh responses to a meal, and that loss of vagal gut-brain signaling dampens HPCd ACh tone.

**Figure 4:**
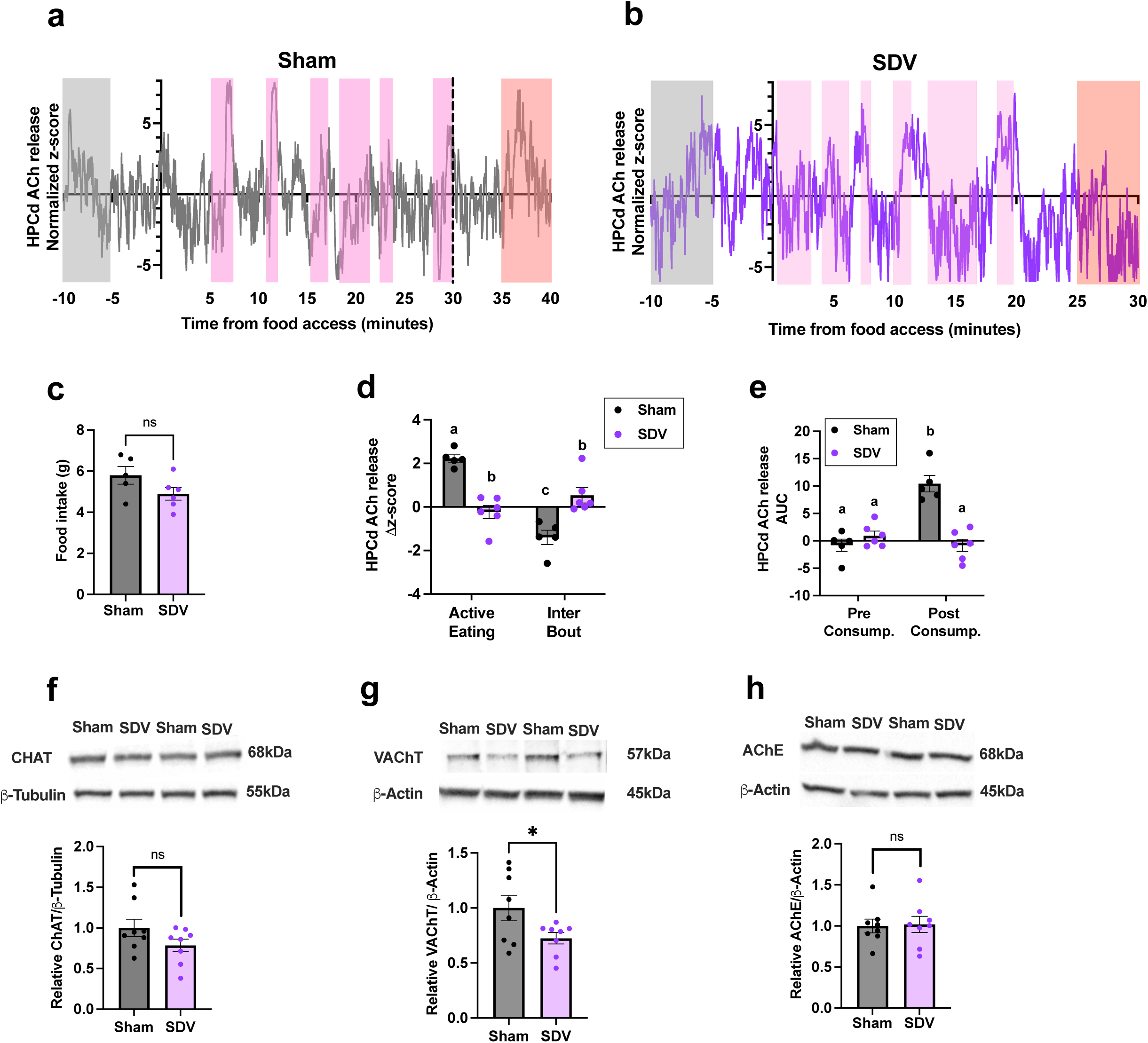
Meal-induced septo-hippocampal cholinergic signaling requires the vagus nerve. A. and. **B.** Representative trace of Sham **(A)** and SDV **(B)** rat HPCd ACh release during consumption of standard chow, time locked to moment of access to food. Pre-consumption period (fasted state) is marked in the gray box, active eating periods [pink boxes; quantified as the Δ in activity between start and end of eating bout], inter bout intervals [white boxes; quantified as the Δ in activity between after the end of an eating bout and before the beginning of the subsequent eating bout], and post-consumption period (satiated state) marked in the red box are represented. **C**. Food intake during fiber photometry recording sessions. **D and E** Fiber photometry recordings of HPCd ACh release during food consumption in animals with either sham surgery (Sham; gray), or SDV (purple). All treatment data containing the same letter above are not significantly different; data were analyzed using two-way ANOVA with multiple comparisons, (*n*=5, Sham; *n*=6, SDV) with **D** depicting change in ACh release during active eating bouts and inter bout intervals for both experimental groups. SDV treatment x eating period interaction \*\*\*\**P<*0.0001. Sham, active eating vs. inter bout \*\*\*\**P*<0.0001; Active eating, Sham vs. SDV \*\*\*\**P*<0.0001; Inter bout, Sham vs. SDV \*\*\**P*=0.0006. **E.** HPCd ACh release AUC quantification during pre-consumption and post consumption periods. SDV treatment x energy state interaction \*\**P=*0.0073. Sham, pre consumption vs. post consumption \*\*\**P*=0.00017; Post consumption, Sham vs. SDV \*\*\**P*=0.0002. **F.** Immunoblot representative images and results for choline acetyltransferase (ChAT) protein levels in the HPCd of rats following sham or SDV surgery (*n*=8, Sham; *n*=8, SDV; paired two-tailed student’s *t*-test, *P*=0.125). **G.** Immunoblot representative images and results for vesicular acetylcholine transporter (VAChT) protein levels in the HPCd of rats following sham or SDV surgery (*n*=8, Sham; *n*=8, SDV; paired two-tailed student’s *t*-test, *P*=0.0154). **H.** Immunoblot representative images and results for acetylcholinesterase (AChE) protein levels in the HPCd of rats following sham or SDV surgery (*n*=8, Sham; *n*=8, SDV; paired two-tailed student’s *t*-test, *P*=0.888). \**P*<0.05. Data are shown as mean ± SEM.

### Western diet consumption blunts meal-induced hippocampal acetylcholine responses and CCK’s satiating effects

Given that Western diet (WD) consumption is associated with both impaired hippocampal-dependent memory and blunted vagal gut-to-brain responses^18,19,21–23,34,35^, we next assessed whether early-life WD diet consumption impacts meal-induced HPCd ACh responses and satiation responses to CCK. Animals were exposed to a cafeteria-style (CAF) WD for 30 days starting from postnatal day (PN) 26 and then switched to a standard chow diet prior to experimental testing from PN 60-85 (Figure 5a). Consistent with previous studies from our laboratory^18,21,36,37^ early life WD exposure did not affect body weights relative to standard chow fed animals (Supp. Figure 5a). Fiber photometry recordings during meal consumption revealed that early life exposure to a WD spared ΔHPCd ACh release elevations during active eating periods (Figure 5e). However, fasted vs satiated analyses revealed that, unlike controls, CAF animals did not experience a sustained elevation in HPCd ACh release after voluntary meal termination (Figure 5f) despite having consumed comparable amounts during the session (Figure 5d). Furthermore, meal period analyses revealed controls but not CAF animals showed elevated magnitude of ΔHPCd ACh release during late vs. early eating bouts (Supp. Figure 5b-e). To examine the long-lasting impacts of our early-life CAF WD model on vagal afferent nerve-mediated satiation signaling, we performed CCK food intake analyses during the healthy diet access period where fasted animals received either a low dose of CCK or vehicle IP, and chow intake was measured 30- and 60-minutes after. Results revealed that IP CCK (0.5μg/kg) significantly reduced food intake at 30-minutes in animals raised on a standard chow diet, whereas CAF animals did not display reductions in food intake from IP CCK treatments at either time-point (Figure 5g). Additionally, automated-home cage food intake measurements revealed that early-life CAF animals consumed significantly more food (chow) on average over a 6-day period than their control counterparts (Figure 5h-i), and that this effect was driven by an increase in the average size of meals collapsed over the 6-day period (Figure 5j-l). In summary, these results reveal that early-life WD exposure blunts vagally-mediated HPCd ACh responses to meal consumption, as well as the satiating response to a vagally-mediated dose of peripheral CCK well into adulthood. Thus, it is plausible that these impairments in vagus-brain signaling underlie hippocampal-dependent memory impairments associated with WD consumption.

**Figure 5:**
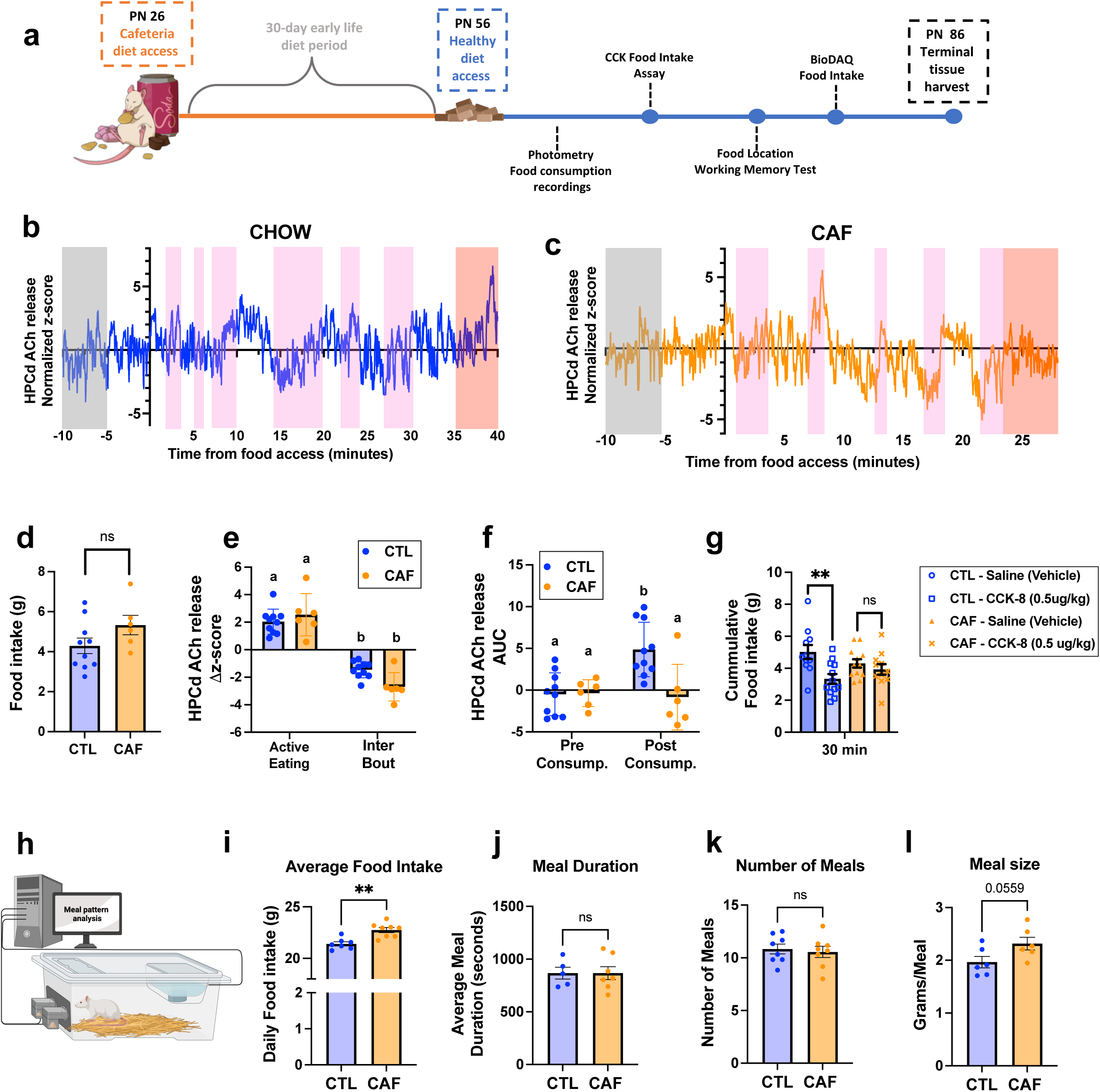
Western diet blunts gut-vagal signaling and eliminates meal-induced septohippocampal cholinergic responses. **A.** Experimental design timeline for fiber photometry and behavioral assessments. **B and C.** Representative trace of chow fed **(B)** and cafeteria (CAF) fed **(C)** rats HPCd ACh release during consumption of standard chow, time locked to moment of access to food. Pre-consumption period (fasted state) is marked in the gray box, active eating periods [pink boxes; quantified as the Δ in activity between start and end of eating bout], inter bout intervals [white boxes; quantified as the Δ in activity between after the end of an eating bout and before the beginning of the subsequent eating bout], and post-consumption period (satiated state) marked in the red box are represented. **D**. Standard chow intake during fiber photometry recording sessions. **E and F** Fiber photometry recordings of HPCd ACh release during food consumption in control (chow fed; CTL) and CAF animals (CTL; blue), or CAF (orange). All treatment data containing the same letter above are not significantly different; data were analyzed using two-way ANOVA with multiple comparisons, (*n*=10, CTL; *n*=6, CAF) with **E.** depicting change in ACh release during active eating bouts and inter bout intervals for both experimental groups. Diet treatment x eating period interaction *P=0.239*. CTL, active eating vs. inter bout \*\*\*\**P*<0.0001; CAF, active eating vs. inter bout \*\*\*\**P*<0.0001; Active eating, CTL vs. CAF *P*=0.635; Inter bout, CTL vs. CAF *P*=0.0593. **F.** HPCd ACh release AUC quantification during pre-consumption and post consumption periods. Diet treatment x energy state interaction \**P=*0.0116. CTL, pre-consumption vs. post-consumption\*\*\**P*=0.0007; Post-consumption, CTL vs. CAF \*\**P*=0.0017. **G.** 30-minute cumulative standard chow intake of CTL and CAF rats after IP saline (vehicle) or CCK (0.5μg/kg). All treatment data containing the same letter above are not significantly different; data were analyzed using two-way ANOVA with multiple comparisons, (*n*=12, CTL; *n*=12, CAF). Diet x IP Treatment interaction *P*=0.1534. CTL, vehicle vs. CCK \*\**P=*0.0063; CAF, vehicle vs. CCK *P*=0.2786; Vehicle, CTL vs. CAF *P*=0.7546; CCK, CTL vs. CAF *P*=0.4754. **H.** Schematic of rats in the BIODAQ system for automated meal pattern analyses. **I.** Average daily food intake over 6-day period in CTL and CAF animals \*\**P*=0.0011. **J.** Average daily meal duration in seconds, *P*=0.9985. **K.** Average daily number of meals, *P*=0.7063. **L.** Average daily meal size expressed as grams per meal, *P*=0.0559. \*\**P*<0.01. Data are shown as mean ± SEM.

### Episodic memory for a meal requires the vagus nerve and septal cholinergic neurons, and is impaired by Western diet consumption

To determine the effects of the experimental models used in this study on meal-associated hippocampal-dependent memory performance; animals were subjected to our “meal place recognition” procedure, which assesses working memory for the spatial location of recent eating. In this procedure hungry rats consumed food in a hidden tunnel in a Barne’s maze, and were then given a second trial shortly thereafter to test for food location memory (Figure 6a). Memory probe test performance results revealed that animals injected with 192IgG saporin in the MS, SDV, or CAF animals had meal location-associated memory impairments, evidenced by their learning index within the probe test being significantly lower than that of their control counterparts (Figure 6b-d). Further, while 192IgG saporin and SDV animals performed significantly worse than control saporin and sham animals, respectively, they also did not perform better than chance. Although CAF animals underperformed relative to their control counterparts, their performance was significantly above chance (Figure 6d), suggesting that early life CAF diet blunts gut-vagal-hippocampus signaling, but to a lesser degree than ablation of the vagus nerve or MS ACh neurons. Together, these data identify memory impairments for the location of a recently consumed meal across three distinct yet related experimental models that disrupt HPCd ACh responses to vagally-mediated GI signals. Further, these findings suggest that vagus-HPC signaling is degraded by early life WD via aberrant septo-hippocampal cholinergic signaling.

**Figure 6:**
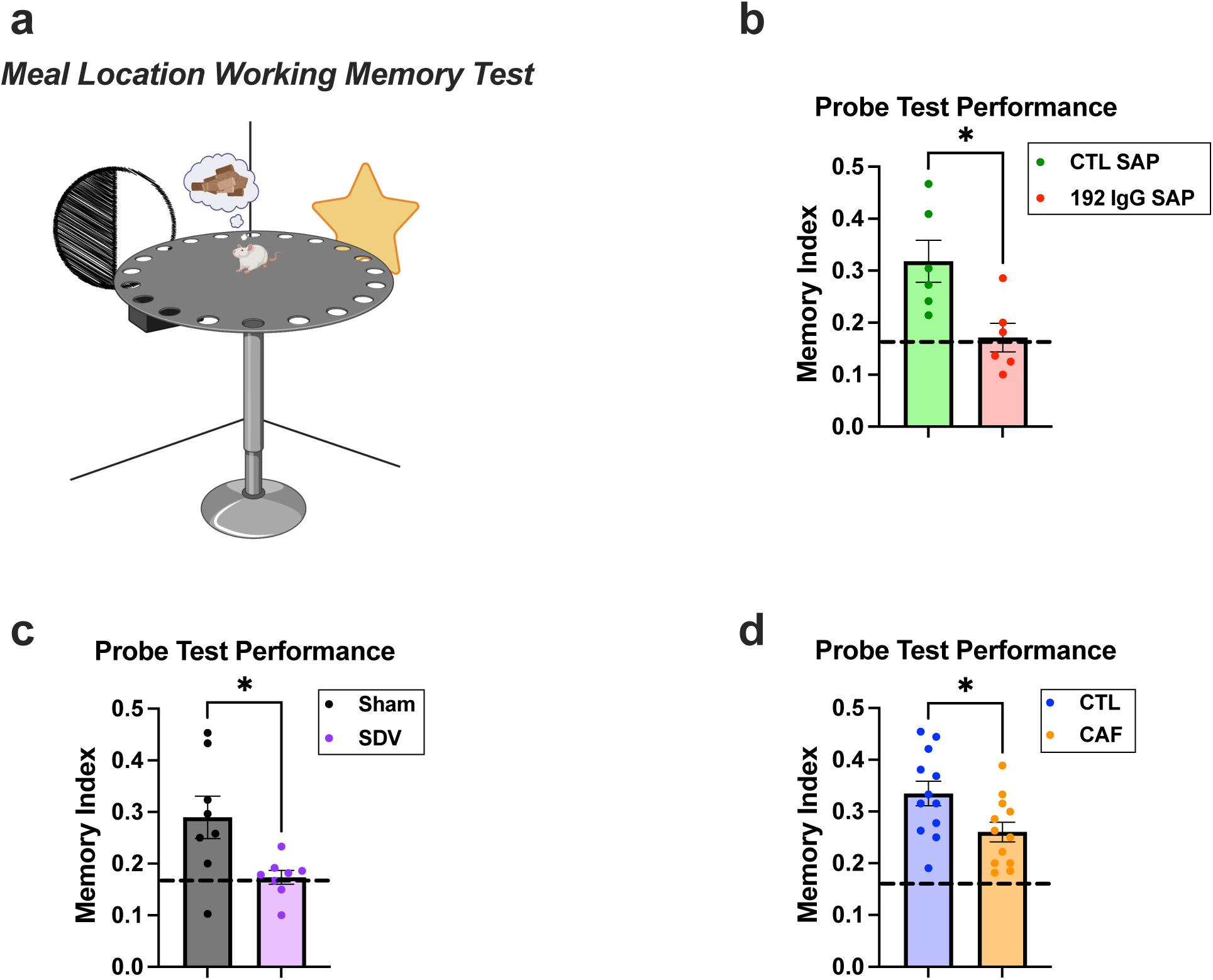
Medial septum cholinergic neuron ablation, subdiaphragmatic vagotomy, and early life Western diet induce meal-associated spatial memory impairments. **A.** Schematic of Meal Location Working Memory Paradigm. **B.** Memory performance during probe test (expressed as learning index) in animals with control saporin in the MS (CTL SAP; green) or 192IgG SAP (red) in the MS; (*n*=6, CTL SAP; *n*=6 192IgG SAP) CTL SAP vs. 192IgG SAP \**P=*0.134; CTL SAP null hypothesis test \**P*=0.0135; 192IgG SAP null hypothesis test *P*=0.8681. **C.** Memory performance during probe test (expressed as memory index) in animals that received either sham surgery (sham; gray) or SDV (purple); (*n*=8, Sham; *n*=8 SDV) Sham vs. SDV \**P=*0.0379; Sham null hypothesis test \**P*=0.0200; SDV null hypothesis test *P*=0.6287. **D.** Memory performance during probe test (expressed as learning index) in chow fed (CTL; blue) animals or CAF fed animals (orange); (*n*=12, CTL; *n*=12, CAF) CTL vs. CAF \**P=*0.0233; CTL null hypothesis test \*\*\*\**P*<0.0001; CAF null hypothesis test \*\*\**P*=0.0005. All treatment comparisons were done using two-tailed paired student’s *t*-test. Null hypothesis testing was done using one-sample *t*- and Wilcoxon test with 0.1667 as the hypothetical null value representing correct hole investigations made by chance. Data are shown as mean ± SEM.

## Methods

### Animals

Male Sprague–Dawley rats (Envigo; 300–400 g; and 70-90 g for CAF diet experiment on arrival) were individually housed with ad libitum access (except where noted) to water and chow (LabDiet 5001, LabDiet, St. Louis, MO) on 12 h:12 h light/dark cycle (lights on at 23:00 h). All procedures involving animals were approved by the University of Southern California Institute of Animal Care and Use Committee.

### Intracranial viral injection and in vivo photometry optic fiber placement

Rats were anesthetized and sedated via intramuscular injection of a ketamine (90.1 mg/kg), xylazine (2.8 mg/kg), and acepromazine (0.72 mg/kg) cocktail. This was followed by a preoperative subcutaneous injection of the analgesic buprenorphine SR (1.3 mg/kg). Once sedated, the surgical site was shaved and disinfected with iodine and ethanol swabbing before the animals were secured in a stereotaxic apparatus. Adeno-associated virus (AAV) encoding iAChSnFR (1 μL; original titer ≥ 7×10¹² vg/mL, diluted 1:2 in artificial cerebrospinal fluid; pAAV.hSynap.iAChSnFR, 137950-AAV1, Addgene, Watertown, MA, USA)—previously validated as a selective in vivo ACh sensor in rodents^38–41^ was injected bilaterally into the dorsal dentate gyrus of the hippocampus (HPCd) using a micro infusion pump (Harvard Apparatus, Cambridge, MA, USA) equipped with a 33-gauge micro syringe injector connected to a PE20 catheter and Hamilton syringe. The injection was delivered at a rate of 5 μL/min, with injectors left in place for an additional 2 minutes to ensure complete infusion. Following viral delivery, fiber optic cannulae (flat 400 μm core, 0.48 numerical aperture, 5 mm; Doric Lenses Inc., Quebec, Canada) were implanted at the same stereotaxic coordinates as the viral injection (A/P: - 3.24; M/L: ±1.80; D/V: -3.50, with bregma as the reference point). The optic fibers were secured to the skull using jeweler’s screws, instant adhesive super glue, and dental cement.

For medial septum cholinergic neuron ablation, the described preoperative methodology was repeated. The antineuronal immunotoxin 192IgG saporin, which has been validated as an agent that selectively ablates neurons that produce acetylcholine^10,42,43^, was injected in the medial septum; 200nL (0.16 μg/μL) were infused in three different coordinates that span the medial septum (A/P_1_: 0.5, M/L_1_: 0.5, D/V_1_: -7.4, A/P_2_: 0.5, M/L_2_: 0, D/V_2_: -6.8, A/P_3_: 0.5, M/L_3_: -0.5, D/V_3_: -7.4, with bregma as the reference point). Rats were allowed to recover after surgery for 2 weeks before experiments proceeded.

### In vivo fiber photometry

Fiber photometry recordings were conducted following previously established methods^21,44,45^. Signal acquisition was performed using the Neurophotometrics fiber photometry system (Neurophotometrics, San Diego, CA) at a sampling frequency of 40 Hz, with alternating excitation wavelengths of 470 nm (ACh-dependent) and 415 nm (ACh-independent). Fluorescence was transmitted through an optical patch cord (Doric Lenses) and directed onto the implanted optic fiber, which in turn relayed neural fluorescence back through the same optic fiber and patch cord to a photoreceiver. Behavioral events, including cues, entries, and eating bouts, were time-stamped using the data acquisition software (Bonsai). To correct for baseline neural activity and motion artifacts, the ACh-independent signal was subtracted from the ACh-dependent signal, and the resulting fluorescence fluctuations were fitted to a biexponential curve. The corrected fluorescence signal was then normalized for each rat by calculating ΔF/F using the average fluorescence signal from the entire recording, followed by conversion to z-scores.

Finally, the normalized signal was aligned to relevant behavioral events, and data extraction was performed using custom MATLAB code.

### Fiber photometry during meal consumption

Prior to the test day, rats underwent an overnight fast with *ad libitum* access to water. Testing occurred in the early nocturnal/dark phase for a 24-hr period without food. On test days, animals were first placed in a neutral context (neutral rat cage in a procedure room with dim lighting) for 15 minutes without food, followed by a 30-minute chow access period (Laboratory Rodent Diet 5001, St. Louis, MO, USA). Afterward, food was removed, and animals remained in the context for an additional 10 minutes. Eating bouts and inter bout intervals were manually time-stamped within the photometry data acquisition stream by an experimenter observing the animals eat on a video monitor system. To assess the effects of physiological HPCd ACh release during refeeding after an overnight fast, in vivo fiber photometry was performed. A patch cord was connected to the implanted optical fiber, and LEDs alternated between 470 nm (ACh-dependent) and 415 nm (ACh-independent) wavelengths throughout the task. The 5-min pre-consumption period for each animal was defined as a 5-min period terminating 5 min before food access. The 5-min post-consumption period was defined as a 5-min period immediately after voluntary meal termination (5 minutes or longer without eating), or the 5-min period beginning 5 min after food removal for animals that did not terminate a meal within the 30-min food access period.

Liquid meal consumption analyses were conducted under similar conditions, except that the consumption tests occurred in operant boxes with lickometers to deliver solutions (Med Associates Inc., Fairfax, VT).

### Subdiaphragmatic vagotomy (SDV)

Animals were habituated to a liquid diet (Research Diets; AIN76A) for five days prior to surgery. Following a 24-hour fast, surgeries were performed under anesthesia with ketamine (90.1 mg/kg), xylazine (2.8 mg/kg), and acepromazine (0.72 mg/kg), along with analgesia provided by Metacam (2 mg/kg). The trunks of the subdiaphragmatic vagus nerve were transected as previously described ^10,46^. A midline abdominal incision was made, after which the stomach was retracted caudally and the liver cranially to expose the esophagus. The dorsal and ventral branches of the vagus nerve were carefully dissected from the esophagus, ligated twice with surgical thread at 1–2 cm intervals, and then cauterized between the ligatures. In sham surgeries, the vagal trunks were exposed but neither ligated nor cauterized. The incision was closed using running sutures for the abdominal wall and stop sutures for the skin. Rats were allowed to recover until their liquid diet intake stabilized (a minimum of one week), after which they were transitioned back to a standard chow diet.

### Dietary model

To simulate a Western “junk food” diet during early life, we implemented a junk food cafeteria-style (CAF) diet, a model previously developed in our laboratory ^21,37^. This diet provided ad libitum access to a high-fat, high-sugar chow (HFHS diet; D12415, Research Diets, New Brunswick, NJ, USA; 20% kcal from protein, 35% kcal from carbohydrates, 45% kcal from fat), along with potato chips (Ruffles Original, Frito Lay, Casa Grande, AZ, USA), chocolate-covered peanut butter cups (Reese’s Minis Unwrapped, The Hershey Company, Hershey, PA, USA), and an 11% weight/volume (w/v) high-fructose corn syrup-55 (HFCS) beverage (Best Flavors, Orange, CA, USA). Each food and drink item were placed in separate receptacles within the home cage. The 11% w/v HFCS concentration was selected to match the sugar content of commonly consumed sugar-sweetened beverages, as previously modeled. CAF-fed rats also had ad libitum access to water. Control (CTL) rats were provided with the same number of food and drink receptacles; however, these contained only standard chow (LabDiet 5001; LabDiet, St.

Louis, MO, USA; 28.5% kcal from protein, 13.5% kcal from fat, 58.0% kcal from carbohydrates) and water. Body weight and food intake—including any spillage collected from cardboard placed beneath the hanging wire cages—were measured three times per week between 8:30 and 10:30 AM, just before the onset of the dark cycle. For the CAF diet model experiments, juvenile rats arrived at the animal facilities on postnatal day (PN) 25. The subsequent day (PN 26), rats were randomly assigned to either the CAF or CTL group (n=12 per group) and given ad libitum access to their respective diet. At PN 56, the CAF group of rats were switched to the ‘healthy’ standard chow diet (LabDiet 5001), the same diet as the CTL group. Experiments with this dietary model were performed after the CAF group had been switched to/acclimated to the ‘healthy’ standard chow diet (PN 63+).

For meal pattern analyses, animals were housed in a BioDAQ food monitoring intake system (Research Diets Inc., New Brunswick, NJ) to which they were habituated to for at least 5 days prior to any recordings. An inter-meal interval (IMI) of 900 s was used to separate individual eating bouts into separate meals. Cumulative chow intake and meal patterns were recorded for 6 consecutive days.

### c-Fos Protein Expression

Surgically treated (MS saporin injections) rats (n = 17) received intraperitoneal (IP) injections of either saline (n = 8) or CCK (CCK-8, 3.0 μg/kg; Bachem, n = 9) 90 minutes prior to perfusion. Tissue was then collected and processed for immunohistochemistry (IHC) as described above. Animals were anesthetized and transcardially perfused with 0.9% saline, followed by 4% paraformaldehyde (PFA) in 0.1 M borate buffer (pH 9.5). Brains were extracted and post-fixed in 12% sucrose in PFA at 4°C for 24 hours. Following fixation, brains were rapidly frozen using dry ice-cooled isopentane and sectioned at 30 μm using a sliding microtome. Tissue sections were stored at –20°C in antifreeze solution until immunohistochemical (IHC) processing. For IHC, all antibody incubations were conducted at 4°C, while washes and other procedural steps were performed at room temperature. Sections were permeabilized with 0.3% Triton X-100 (Sigma, Cat #: X100-500ML) for 30 minutes, then blocked for 10 minutes with 2% normal donkey serum (Jackson ImmunoResearch, Cat #: 017-000-121). Primary antibodies were diluted in KPBS containing 2% normal donkey serum and applied overnight (∼18 hours). Following primary incubation, sections were washed in KPBS (6 changes, 10 minutes each). Secondary antibodies conjugated to fluorophores were also diluted in KPBS with 2% donkey serum and applied overnight. After secondary incubation, sections were again washed in KPBS (2 changes, 2 minutes each), then mounted on glass slides, air-dried, and coverslipped using 50% glycerol in KPBS as the mounting medium. Detection of c-Fos was performed using a rabbit anti-c-Fos primary antibody (1:500, Cell Signaling, Catalog No. 2250S), followed by a donkey anti-rabbit IgG-AlexaFluor AF488 secondary antibody (1:500, Jackson Immunoresearch, RRID: AB_2340619). Representative images and quantification of c-Fos protein expression in CA3d and DG regions were obtained from both saline- and CCK-injected animals, with analysis confined to Swanson Atlas levels 28–30. Fluorescent images were captured using a Nikon 80i microscope equipped with a DS-Qi1 camera (1280 × 1024 resolution, 1.45 megapixels) under epifluorescent illumination. Image acquisition was performed using Nikon Elements BR software.

### Western Blotting

Dorsal HPC tissue punches from the SDV experiment were analyzed for levels of markers of cholinergic tone (choline acetyltransferase [ChAT], vesicular acetylcholine transporter [VAChT], and acetylcholinesterase [AChE]). Proteins in brain lysates (25 μg per sample/well; determined via the reducing agent and detergent compatible [RC DC^TM^] protein assay, Bio-Rad Laboratories, Inc.) were separated using sodium dodecyl sulfate polyacrylamide gel electrophoresis, transferred onto poly-vinylidene difluoride membranes, and subjected to enhanced chemiluminescence for immunodetection analysis (Chemidoc XRS, BioRad). A rabbit polyclonal anti-choline acetyltransferase antibody (1:100; Sigma-Aldrich, catalog # AB143) was used to determine the concentration of ChAT relative to a load control signal detected by a rabbit anti-β-tubulin antibody (1:1000; Cell Signaling Technology, 9F3 rabbit mAb, catalog # 2128). A rabbit polyclonal anti-vesicular ACh antibody (1:1000; Sigma-Aldrich, catalog # ABN100) was used to determine the concentration of VAChT relative to a load control signal detected by a rabbit anti-β-actin antibody (1:1000; Abcam, catalog # ab8227). A rabbit recombinant anti-acetylcholinesterase antibody (1:1000; Abcam, catalog # ab183591) was used to determine the concentration of AChE relative to a loading control signal detected by a rabbit anti-β-actin antibody (1:1000; Santa Cruz Biotechnology, catalog # NB600-503). Goat anti-rabbit-IgG-horseradish peroxidase (HRP)-linked antibody (1:5000; Cell Signaling Technology, catalog # 7074) was used as the secondary antibody. Blots were quantified with densitometry analysis using ImageJ as previously reported ^10,21^.

### Meal Location Memory Procedure

Meal location memory was assessed using an elevated circular Barnes maze (Diameter: 122 cm, Height: 140 cm) with 18 evenly spaced holes around its circumference. A hidden black escape box (38.73 cm × 11.43 cm × 7.62 cm) was placed beneath one of the holes. To provide visuospatial cues, four distinct markers (black and white stripes, a white circle, a red triangle, a stuffed unicorn, and a collection of irregular shapes) were positioned on the surrounding walls. One day after habituation, spatial working memory training commenced. Rats underwent mild food restriction, with food removed one hour before the onset of their dark cycle. In the maze, they relied on visuospatial cues to locate the escape box, which contained five sucrose pellets. Each rat completed two trials per day for five consecutive training days, with a two-minute interval between trials during which the maze was cleaned and rotated. The escape box remained in the same location for both trials on a given training day but was repositioned at the start of the first trial each subsequent day.

Errors were recorded as investigations of holes without the escape box or the two holes adjacent to the escape box, while meal location working memory was quantified as the difference in errors between trial 2 and trial 1 on each training day. These values were then averaged across the five training days. A memory probe test was conducted in which animals underwent testing identical to training, however, no escape box was present. Performance (memory index) in the probe test was quantified by calculating the number of correct investigations divided by the total number of investigations executed during the second trial.

### CCK-8 and Amylin treatment with fiber photometry

Two days before test day, animals were habituated to an intraperitoneal (IP) injection (saline) while being connected to the fiber photometry patch cord (without recording). On test day, animals underwent an overnight fast with *ad libitum* access to water, and then they were placed in a neutral cage for 15 minutes without intervention, after which they received IP injections of either CCK-8 (3μg/kg; Bachem), Amylin (50 μg/kg; Calbiochem), or their corresponding vehicle (sterile saline). To assess the effects of IP administration of these treatments on HPCd ACh release, we performed fiber photometry and recorded from the 15-minute baseline period to 40 minutes after IP administration.

### Statistical analyses

Statistical analyses were performed using GraphPad Prism 9.0 software (GraphPad Software Inc., San Diego, CA, USA) and Microsoft Excel V16.66.1. Data are expressed as mean ± SEM. Statistical details can be found in the figure legends and *n*’s refer to the number of animals for each condition. Differences were considered statistically significant at *P* < 0.05.A two-tailed paired Student’s *t*-test was used to compare within-subject HPCd Ach release for active eating and inter bout periods, as well as pre-meal and post-meal periods during the standard consumption of standard chow, 11% sucrose, 4.5% corn oil, 0.2% saccharin, and 0.2% saccharin + 2% sucrose. Paired Student’s *t-*test was also used to compare within-subject training and probe test data for the meal location working memory task.

A two-tailed unpaired Student’s *t*-test was used to compare the consumption of 11% sucrose to 4.5% corn oil, and 0.2% saccharin + 2% sucrose, as well as 0.2% saccharin + 2% sucrose to 4.5% corn oil, and 2% saccharin between groups. Two-tailed unpaired Student’s *t*-test was also used to compare standard laboratory chow consumption during fiber photometry meal consumption recordings between CTL SAP and 192IgG SAP, Sham and SDV (Western blot, meal location recognition), as well as CTL and CAF group (meal location recognition and meal patterns). Outliers were identified using the Grubb’s test for outliers post-hoc at significance level of alpha = 0.05. For all experiments, assumptions of normality, homogeneity of variance (HOV), and independence were met where required.

All comparisons of the effects of either 192IgG SAP, SDV, and CAF, c-fos expression, body weight, and HPCd ACh release responses were analyzed using a two-way ANOVA with Bonferroni’s post hoc test for multiple comparisons. For all statistical tests, the α level for significance was 0.05.

## Discussion

Despite recent findings implicating the vagus nerve in the modulation of cognitive processes ^5,10^, the neural substrates driving these phenomena, particularly promoting HPC-dependent memory function, remain poorly understood. Our findings establish a novel gut-brain substrate through which nutrient-induced activation of gastrointestinal VAN engages HPCd ACh signaling via MS cholinergic neurons. This neuronal pathway is responsive to vagally-transmitted satiation signals, including CCK and physiological meal consumption, and is disrupted by early-life WD exposure. These data offer fundamental mechanistic insights into how peri-prandial physiological signals shape HPC activity and thus contribute to our understanding of how interoceptive signals from the gut affect neurocognitive processes.

Our results reveal that peripheral administration of CCK increases HPCd c-Fos expression and ACh signaling tone, effects that are abolished by selective ablation of MS cholinergic neurons (Figure 1). This provides causal evidence that MS cholinergic neurons serve as a key relay for conveying vagal information to the HPC, particularly the dorsal dentate gyrus (DG) and CA3 subfields. While the MS has long been known to regulate hippocampal theta rhythm and memory encoding through its cholinergic output ^47,48^, our work identifies an upstream GI influence on this circuit, mediated through GI VAN.

Impairments in meal-location memory were observed across all experimental models that disrupt vagal-ACh-HPC communication—SDV, MS cholinergic ablation (192 IgG SAP) and early-life WD exposure. Given the established role of HPCd ACh transmission in memory encoding ^14–16^, and that WD-induced HPC impairments occur through dysregulation of ACh transmission ^21^, our novel data showing parallel disruptions in meal-associated HPCd ACh release in SDV and CAF animals (Figure 4 & 5) provide evidence that a junk food diet and overall loss of integrity of VAN signaling yields cognitive impairments via disruptions in vagal-ACh-HPC signaling. However, while CAF animals exhibited milder memory deficits compared to vagotomized or MS ACh neuron ablated rats, this may reflect the persistence of other HPC inputs and/or compensatory mechanisms. Nonetheless, the consistency across models underscores the critical role of the vagal-ACh-HPC axis in encoding episodic memories related to eating events.

Fiber photometry recordings reveal robust increases in HPC ACh release during active voluntary food consumption, particularly later in the meal and during the post-meal satiety phase (Figure 2 & Supp Figure 2c-d). Such results demonstrate that naturalistic eating modulates the release of HPCd ACh—a neuromodulator classically associated with attention, learning, and synaptic plasticity ^14,49,50^, during consumption and the transition to satiation and meal termination. The temporally-specific increase in cholinergic tone particularly during later meal phases suggests that hippocampal processing may help encode contextual cues linked to nutrient consumption, thus contributing to episodic memory formation regarding the context of the meal experience. These findings resonate and are consistent with studies demonstrating that the HPC encodes interoceptive states such as hunger and satiety, and HPC activity is functionally modulated by feeding-related cues (e.g., gut hormones) ^51,52^. Notably, the effects of CCK on HPC activation were selective to cholinergic MS neurons were not observed with another feeding-related peptide, amylin, that exerts its anorectic effects independently of vagal transmission ^30^. This specificity highlights vagally-mediated physiological nutrient signaling as a key regulator of HPC neurophysiology.

In dissecting the drivers of meal-induced ACh release, we show that caloric nutrients (sucrose and corn oil) elicit significant HPCd ACh responses, while non- and low-nutritive sweeteners (saccharin, or saccharin + 2% sucrose) do not, despite their palatability and higher ingestion volumes in the latter condition (Figure 3). This nutrient-dependence suggests that HPCd cholinergic activation requires post-ingestive signaling—likely involving gastrointestinal nutrient sensing mechanisms (e.g., nutrient receptors, enteroendocrine cells) that act upstream of vagal afferents ^26,53^. Furthermore, previous work has shown that nutrient sensing in the gut engages vagal circuits to drive behavior and affect central nervous system function ^31,54^. Our data expands this concept by identifying HPC cholinergic tone as a sensitive index of gut-nutrient detection and by implicating this system in the encoding of meal-related memory processes.

We provide direct evidence that SDV abolishes the HPCd ACh responses elicited by nutrient ingestion without altering total caloric intake (Figure 4a-e). Furthermore, vagotomy reduces HPCd vesicular ACh transporter (VAChT) expression, suggesting that chronic vagal disconnection impairs cholinergic transmission capacity, and this happens without compensatory changes in choline-acetyl transferase (ChAT) or acetylcholinesterase (AChE) expression (Figure 4f-h). These findings align with accumulating evidence that vagal input modulates HPC-dependent cognitive processes ^10,55,56^. Importantly, our data link these observations to nutrient-induced changes in cholinergic signaling and identify a mechanism through which gut-brain disconnection can compromise memory function, particularly for meal-associated events. These responses were eliminated by MS cholinergic ablation and by SDV, indicating that both intact VAN and septo-hippocampal cholinergic projections are essential for this nutrient-stimulated HPCd ACh activation.

Consistent with previous findings^17–21,34,37^, early-life consumption of a WD impaired hippocampal-dependent memory. Building on this work, here we show that early-life WD also abolished sustained elevations in post-meal HPC ACh signaling (Figure 5b-f & Figure 6d). While ACh levels rose during active consumption bouts in CAF animals, unlike SDV and MS ACh neuron-ablated rats, the absence of a postprandial ACh elevation suggests that WD exposure altered nutrient-induced satiation signaling later in life. These neurophysiological deficits corresponded with blunted anorexigenic responses to peripheral CCK administration and increased average spontaneous meal size (Figure 5g-l), suggesting a functional disconnection between vagal satiation signals and hippocampal memory systems following early-life WD. Mechanistically, these impairments may stem from WD-induced gut barrier dysfunction, microbiome shifts, and/or VAN remodeling^18,57^ , which have been associated with diminished gut-to-hindbrain communication.

Chronic reductions in ACh signaling markers from early life CAF diet (e.g., VAChT, parallel to those seen in SDV here), are linked with disruptions in HPCd ACh signaling during object-context novelty recognition ^21^. More specifically, when control animals investigated an object that is novel to a specific context, HPCd ACh release increased significantly. This effect, however, was absent in CAF animals who were also impaired behaviorally in the memory task. However, memory performance was rescued through HPCd infusions of a cholinergic (ACh α7 nicotinic) receptor agonist ^21^ during the memory encoding phase of the memory task. Therefore, given the current understanding of how ACh promotes memory encoding processes in the HPC via α7 nicotinic receptor activation ^58,59^, meal-induced HPCd ACh elevations are a likely mechanism that evolved for the purpose of assisting an animals’ capacity to encode the context and environment of when and where food was acquired and consumed for more efficient subsequent foraging sessions. Another possible mechanism through which diet-induced vagal-ACh-HPC disruption causes long-lasting impairments is through non-neuronal mechanisms, as it has been established that ACh has anti-inflammatory effects in the HPC by acting on α7 nicotinic receptors on microglia ^60,61^ and astrocytes ^62^.

The work presented here offers a basic science understanding of how signals from the gut modulate the septo-hippocampal memory system, and how it responds in the presence of environmental insults. These results call for clinical research endeavors to investigate this system in the context of neurodegenerative disorders such as Alzheimer’s Disease (AD) given the established literature showing that AD is characterized by deterioration of the HPC cholinergic signaling ^63^. Taken together with recent emerging findings, collective results demonstrate that dietary insults yield HPC ACh dysregulation parallel to those seen in AD and supports the notion that an obesogenic early-life diet contributes to the development of AD ^64–66^. Consistent with this framework, a population-based cohort study of patients that received an SDV as a form of treatment for gastric ulcers in Taiwan demonstrated a significantly higher risk of developing dementia in SDV individuals vs. control matched individuals ^67^. These findings further highlight the translational importance of the vagus-ACh-HPC circuitry studied here in the context of diet-induced cognitive impairments and AD etiology, and aligns with the existing literature demonstrating the potential benefits of vagus nerve stimulation on memory function ^5,8,68,69^. Therapeutic strategies aimed at restoring vagal tone or enhancing septo-hippocampal cholinergic signaling may thus offer novel avenues for treating obesity-, dietary-, environmental-, or genetic-mediated memory impairments.

---

**Supplementary Figure 1:**
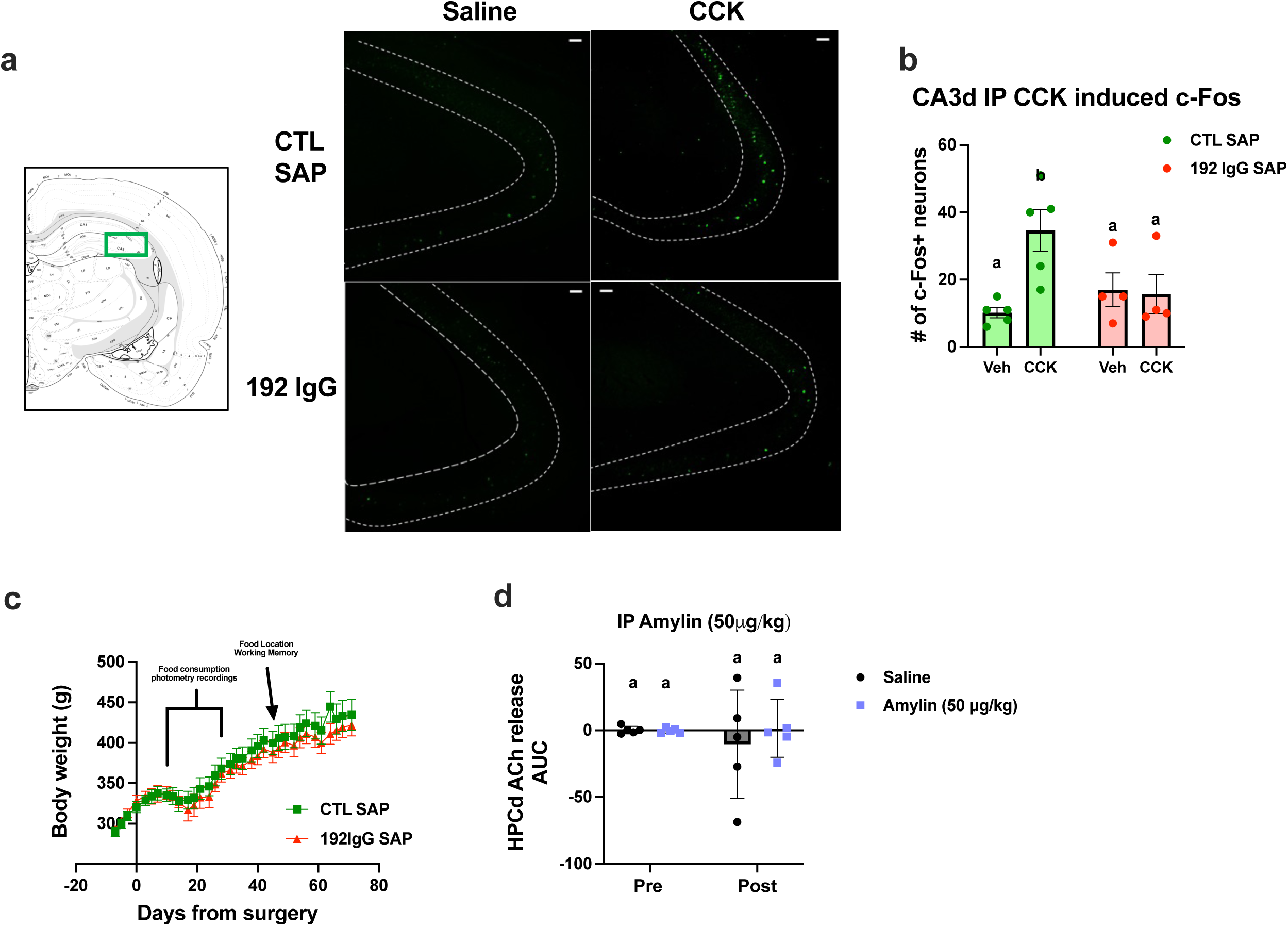
A. Schematic and representative images of dorsal CA3 subregion immunoreactive c-Fos+ neurons in control MS saporin and 192IgG saporin treated animals given IP CCK (3μg/kg) or vehicle (saline). Scale bar = 30 microns. **B.** CA3d c-Fos+ neurons quantification. (*n* = 5 Vehicle-CTL SAP; *n* = 4 CCK-CTL SAP; *n* = 4 Vehicle-192IgG sap; *n* = 4 CCK-192IgG sap) Data were analyzed using two-way ANOVA with multiple comparisons. Treatment data with the same letter above are not significantly different. IP Treatment x saporin interaction \**P =* 0.0209, CTL SAP, Veh vs CCK \*\**P =* 0.0046; CCK treatment, CTL SAP vs. 192IgG SAP \**P =* 0.0342. **C.** Body weight readings of MS CTL SAP (green) and 192IgG SAP (red) animals. Data were analyzed using two-way ANOVA with multiple comparisons; saporin treatment x time interaction *P*=0.6506. **D.** HPCd ACh release AUC quantification in response to IP vehicle and Amylin (50μg/kg) administration. Treatment data with the same letter above are not significantly different. Data were analyzed using two-way ANOVA with multiple comparisons; IP Treatment x time *P =* 0.677; Vehicle, Pre vs. Post *P >* 0.999; Amylin (50μg/kg) P *>* 0.999; Post injection, Vehicle vs. Amylin (50μg/kg) *P* > 0.999. Data are shown as mean ± SEM; \*\*\*\**P*<0.0001, \*\**P<0.01*.

**Supplementary Figure 2:**
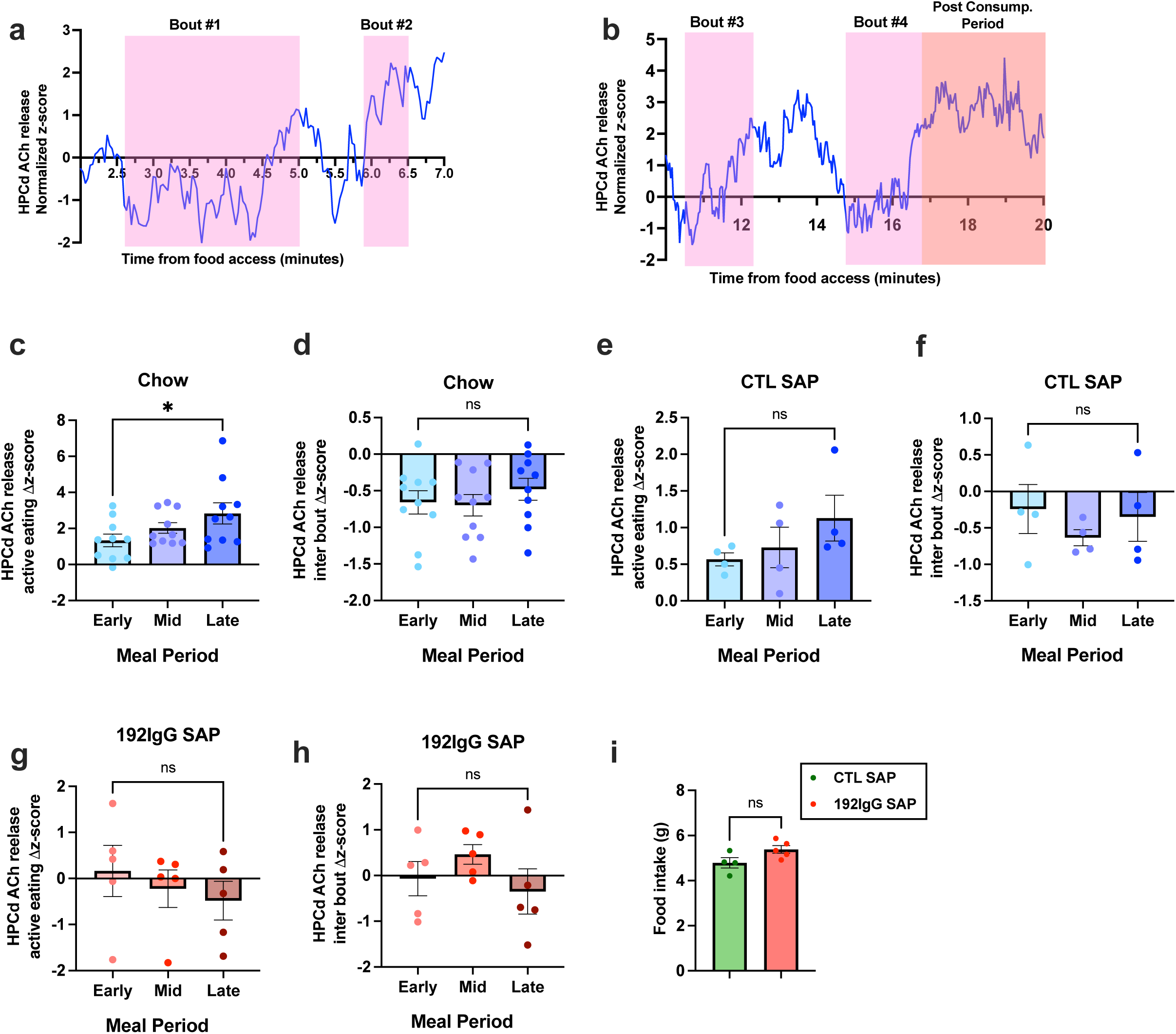
A. Zoomed in graphs of representative trace in main Figure 2B, with **A** depicting the first and second bouts in the session, and **B.** depicting the third, fourth, and post consumption period of the session. **C.** Change in HPCd ACh release during active eating bouts, separated by tertiles of order taken (early, middle, and late bouts). Data were analyzed using two-tailed student’s *t*-test; Early bouts vs. Late bouts \**P* = 0.0289. **D.** Change in HPCd ACh release during inter bout intervals, separated by tertiles of order taken (early, middle, and late intervals). Data were analyzed using two-tailed student’s *t*-test; Early intervals vs. Late intervals *P* = 0.5071. **E.** Change in HPCd ACh release in MS CTL SAP animals during active eating bouts, separated by tertiles of order taken (early, middle, and late bouts). **F.** Change in HPCd ACh release in MS CTL SAP animals during inter bout intervals, separated by tertiles of order taken (early, middle, and late intervals). Data were analyzed using two-tailed student’s *t*-test; Early intervals vs. Late intervals *P* = 0.6004. **G.** Change in HPCd ACh release in MS 192IgG SAP animals during active eating bouts, separated by tertiles of order taken (early, middle, and late bouts). Data were analyzed using two-tailed student’s *t*-test; Early bouts vs. Late bouts *P* = 0.5072. **H.** Change in HPCd ACh release in MS 192IgG SAP animals during inter bout intervals, separated by tertiles of order taken (early, middle, and late intervals). Data were analyzed using two-tailed student’s *t*-test; Early intervals vs. Late intervals *P* = 0.7457. **I.** Standard chow intake during fiber photometry recordings between MS CTL SAP and 192IgG SAP; *P* = 0.0688. Data are shown as mean ± SEM; \**P<0.05*.

**Supplementary Figure 3:**
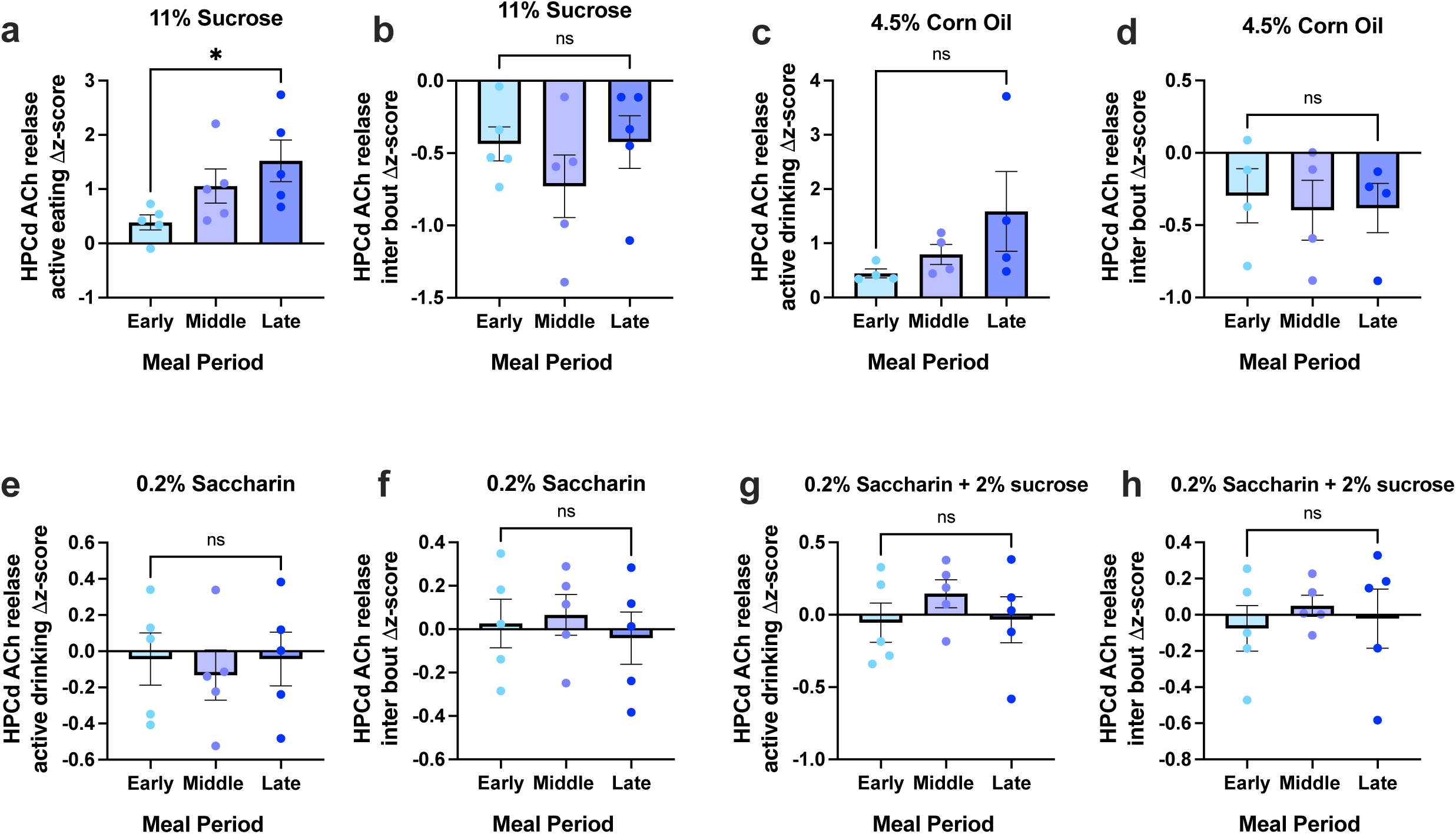
A. Change in HPCd ACh release during 11% sucrose active eating bouts, separated by tertiles of order taken (early, middle, and late bouts). Data were analyzed using two-tailed student’s *t*-test; Early bouts vs. Late bouts \**P* = 0.0470. **B.** Change in HPCd ACh release during 11% sucrose inter bout intervals, separated by tertiles of order taken (early, middle, and late intervals). Data were analyzed using two-tailed student’s *t*-test; Early intervals vs. Late intervals *P* = 0.9107. **C.** Change in HPCd ACh release during 4.5% corn oil active eating bouts, separated by tertiles of order taken (early, middle, and late bouts). Data were analyzed using two-tailed student’s *t*-test; Early bouts vs. Late bouts *P* = 0.2327. **D.** Change in HPCd ACh release during 4.5% corn oil inter bout intervals, separated by tertiles of order taken (early, middle, and late intervals). Data were analyzed using two-tailed student’s *t*-test; Early intervals vs. Late intervals *P* = 0.7290. **E.** Change in HPCd ACh release during 0.2% saccharin active eating bouts, separated by tertiles of order taken (early, middle, and late bouts). Data were analyzed using two-tailed student’s *t*-test; Early bouts vs. Late bouts *P* = 0.9983. **F.** Change in HPCd ACh release during 0.2% saccharin inter bout intervals, separated by tertiles of order taken (early, middle, and late intervals). Data were analyzed using two-tailed student’s *t*-test; Early intervals vs. Late intervals *P* = 0.7039. **G.** Change in HPCd ACh release during 0.2% saccharin + 2% sucrose active eating bouts, separated by tertiles of order taken (early, middle, and late bouts). Data were analyzed using two-tailed student’s *t*-test; Early bouts vs. Late bouts *P* = 0.9450. **H.** Change in HPCd ACh release during 0.2% saccharin + 2% sucrose inter bout intervals, separated by tertiles of order taken (early, middle, and late intervals). Data were analyzed using two-tailed student’s *t*-test; Early intervals vs. Late intervals *P* = 0.8216. Data are shown as mean ± SEM; \**P<0.05*.

**Supplementary Figure 4:**
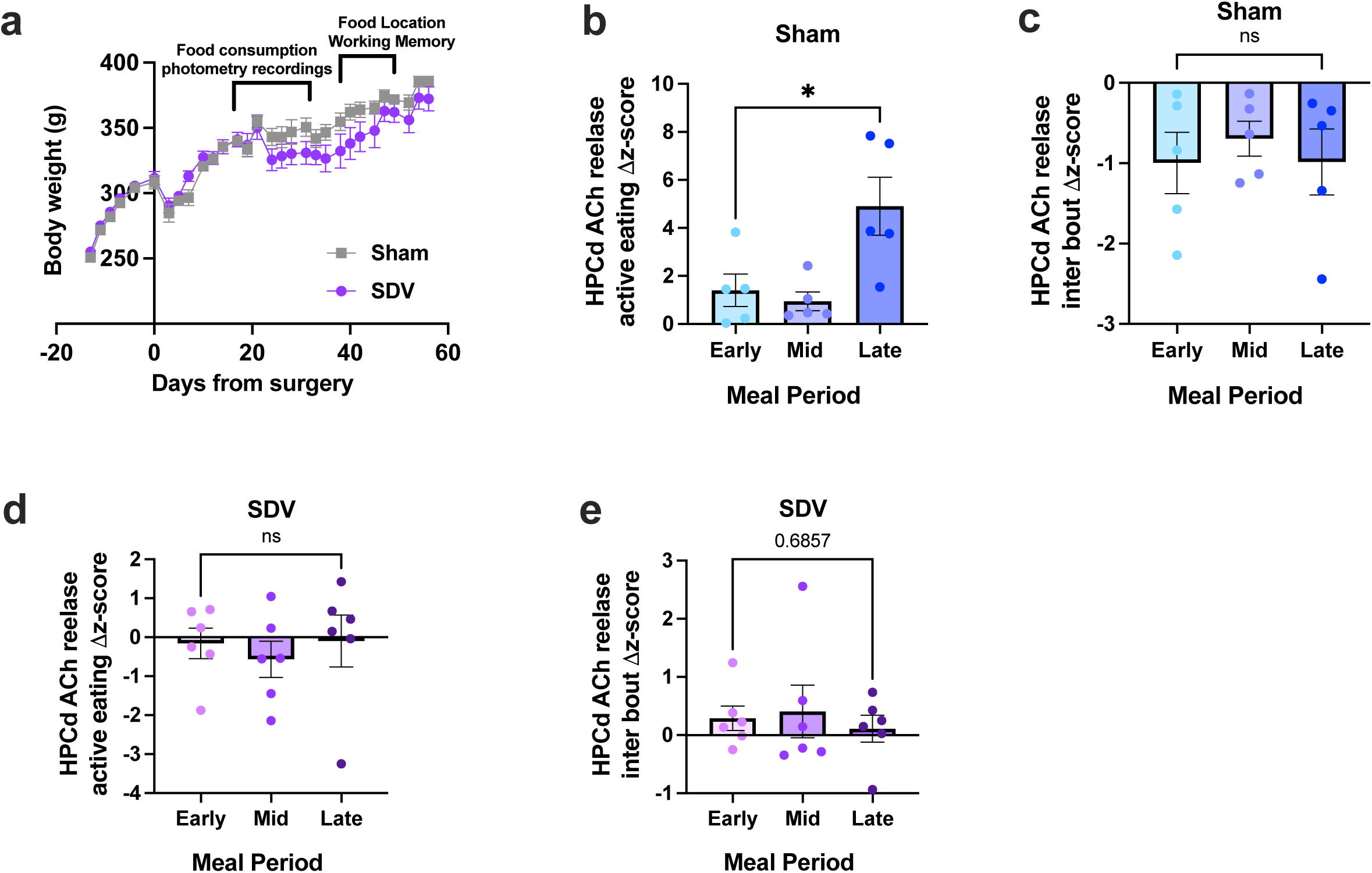
A. Body weight readings of animals that received sham surgery (gray) and SDV (purple) animals. Data were analyzed using two-way ANOVA with multiple comparisons; SDV treatment x time interaction \*\**P*=0.0081. **B.** Change in HPCd ACh release in sham surgery animals during active eating bouts, separated by tertiles of order taken (early, middle, and late bouts). Data were analyzed using two-tailed student’s *t*-test; Early bouts vs. Late bouts *P* = 0.0493. **C.** Change in HPCd ACh release in sham surgery animals during inter bout intervals, separated by tertiles of order taken (early, middle, and late intervals). Data were analyzed using two-tailed student’s *t*-test; Early intervals vs. Late intervals *P* = 0.9725. **D.** Change in HPCd ACh release in SDV surgery animals during active eating bouts, separated by tertiles of order taken (early, middle, and late bouts). Data were analyzed using two-tailed student’s *t*-test; Early bouts vs. Late bouts *P* = 0.9514. **E.** Change in HPCd ACh release in SDV surgery animals during inter bout intervals, separated by tertiles of order taken (early, middle, and late intervals). Data were analyzed using two-tailed student’s *t*-test; Early intervals vs. Late intervals *P* = 0.6857. Data are shown as mean ± SEM; \*\**P<0.05*.

**Supplementary Figure 5:**
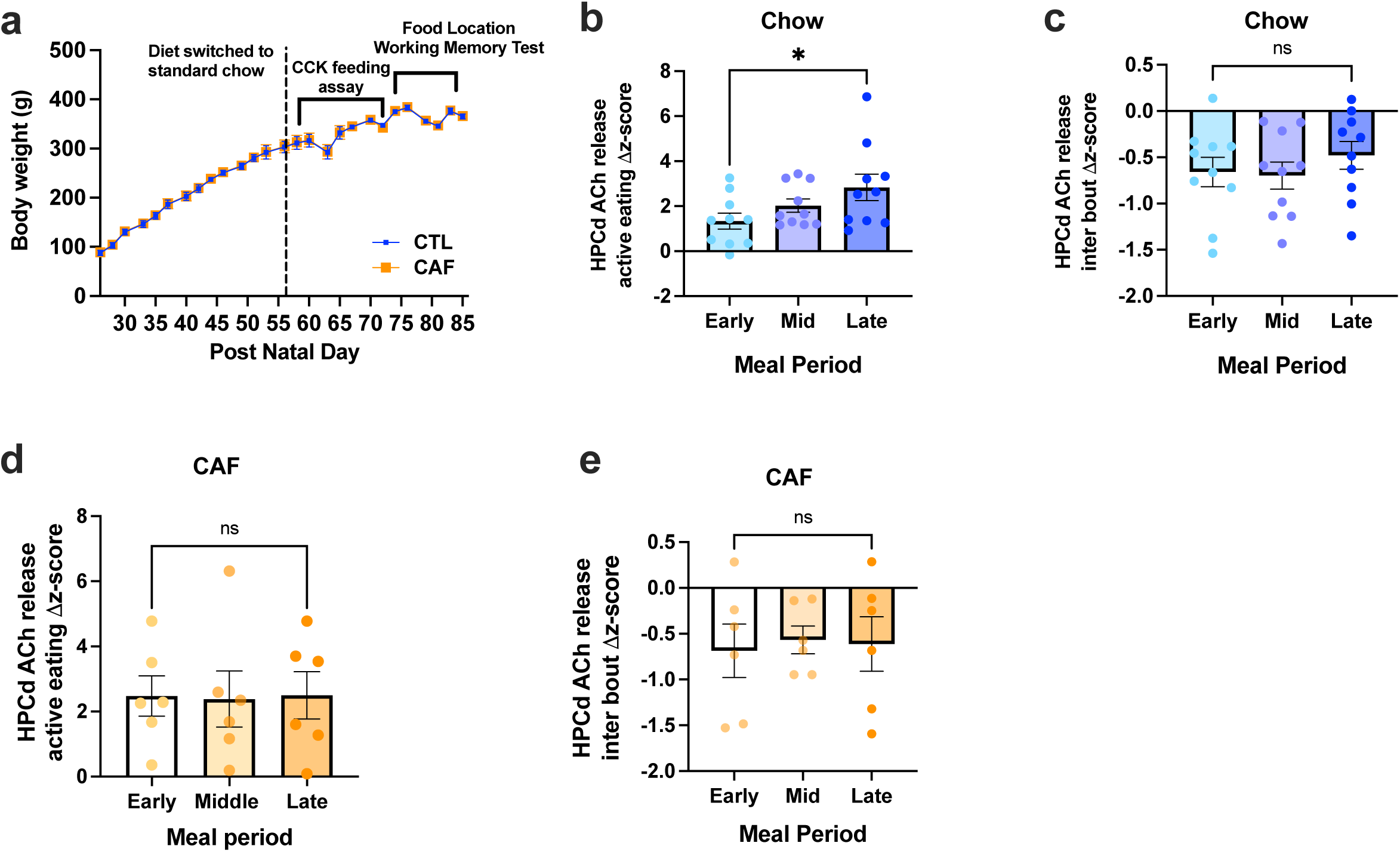
A. Body weight readings of animals that received standard laboratory chow diet (blue) and CAF diet (orange). Data were analyzed using two-way ANOVA with multiple comparisons; Diet treatment x time interaction *P*=0.881. **B.** Change in HPCd ACh release in chow fed animals during active eating bouts, separated by tertiles of order taken (early, middle, and late bouts). Data were analyzed using two-tailed student’s *t*-test; Early bouts vs. Late bouts *P* = 0.0289. **C.** Change in HPCd ACh release in chow fed animals during inter bout intervals, separated by tertiles of order taken (early, middle, and late intervals). Data were analyzed using two-tailed student’s *t*-test; Early intervals vs. Late intervals *P* = 0.5071. **D.** Change in HPCd ACh release in CAF diet fed animals during active eating bouts, separated by tertiles of order taken (early, middle, and late bouts). Data were analyzed using two-tailed student’s *t*-test; Early bouts vs. Late bouts *P* = 0.9811. **E.** Change in HPCd ACh release in CAF diet fed animals during inter bout intervals, separated by tertiles of order taken (early, middle, and late intervals). Data were analyzed using two-tailed student’s *t*-test; Early intervals vs. Late intervals *P* = 0.9025. Data are shown as mean ± SEM; \**P<0.05*.

## Notes

### Competing Interest Statement

The authors have declared no competing interest.

### Funding

- National Institute of Diabetes and Digestive and Kidney Diseases: R01DK104897, R01DK123423, F31DK137484, F31DK138777
- National Institute on Aging: F32AG077932
- Alzheimer's Association: AARFD-22-972811

